# Model evolution in SARS-CoV-2 spike protein sequences using a generative neural network

**DOI:** 10.1101/2022.04.12.487999

**Authors:** Anup Kumar

## Abstract

Modelling evolutionary elements inherent in protein sequences, emerging from one clade into another of the SARS-CoV-2 virus, would provide insights to augment our understanding of its impact on public health and may help in formulating better strategies to contain its spread. Deep learning methods have been used to model protein sequences for SARS-CoV-2 viruses. A few significant drawbacks in these studies include being deficient in modelling end-to-end protein sequences, modelling only those genomic positions that show high activity and upsampling the number of sequences at each genomic position for balancing the frequency of mutations. To mitigate such drawbacks, the current approach uses a generative model, an encoder-decoder neural network, to learn the natural progression of spike protein sequences through adjacent clades of the phylogenetic tree of Nextstrain clades. Encoder transforms a set of spike protein sequences from the source clade (20A) into its latent representation. Decoder uses the latent representation, along with Gaussian distributed noise, to generate a different set of protein sequences that are closer to the target clade (20B). The source and target clades are adjacent nodes in the phylogenetic tree of different evolving clades of the SARS-CoV-2 virus. Sequences of amino acids are generated, for the entire length, at each genomic position using the latent representation of the amino acid generated at a previous step. Using trained models, protein sequences from the source clade are used to generate sequences that form a collection of evolved sequences belonging to all children clades of the source clade. A comparison of this predicted evolution (between source and generated sequences) of proteins with the true evolution (between source and target sequences) shows a high pearson correlation (> 0.7). Moreover, the distribution of the frequencies of substitutions per genomic position, including high- and low-frequency positions, in source-target sequences and source-generated sequences exhibit a high resemblance (pearson correlation > 0.7). In addition, the model partially predicts a few substitutions at specific genomic positions for the sequences of unseen clades (20J (Gamma)) where they show little activity during training. These outcomes show the potential of this approach in learning the latent mechanism of evolution of SARS-CoV-2 viral sequences.

**Codebase:** https://github.com/anuprulez/clade_prediction

## Introduction

Mutations in genomic sequences of viruses are a natural phenomenon that arises as a result of the replication process. They tend to make viruses more compatible with their external surroundings but most of them largely remain silent. However, a few of them may cause severe diseases when they tend to make it favourable for the viruses to evade the immune system of human beings. The adverse effects of such malicious mutations are unknown and pose a severe risk to public health. To get insights into the future space of these genomic sequences, computational methods should be developed to garner knowledge about its evolutionary path. One of the approaches would be to learn the mutations present in genomic sequences at different stages of evolution in the past and to generate sequences yet to be evolved using the learned knowledge. These generated sequences may give useful insights into the mutations that are biologically significant. The impact of such mutations, which are yet to occur, can be further analysed.

### A. SARS-CoV-2

The entire world has been overwhelmed by COVID-19, caused by the SARS-CoV-2 virus, since early 2020. The spread of the virus has caused a pandemic that has adversely impacted the health of scores of human beings in almost every nation. Its impact necessitated the implementation of severe measures, supported by scientific evidence, to minimize the risk, understand the nature of the virus and the disease and protect human lives. Deeper scientific studies have helped sequence the virus and find the proteins associated with the viral genome. Moreover, the evolution of the virus through time has been categorised into multiple categories, also known as Nextstrain (2) clades. A few examples of such clades are 19A, 20B, 21K and many more. In general, a clade is a nomenclature system used by Nextstrain to organize the evolution of several viruses such as Influenza, Enterovirus and SARS-CoV-2 and these clades have signature mutations. In SARS-CoV-2, for example, clade 20B/S has V1122L substitution, clade 21K (Omicron) has many mutations such as A67V substitution, H69-deletion and many more (3). Clades have been arranged in a phylogenetic tree showing the emergence of the SARS-CoV-2 virus since early 2020 (Figure 1), moving from left to right. The genome length of the SARS-CoV-2 virus is around 30 kb that encodes several proteins - 16 non-structural (NSP 1-16) that are involved with tasks such as replication and transcription and 4 structural (spike (S), envelope (E), membrane (M) and nucleocapsid (N)) that are responsible for maintaining the structure of the virus (4).

**Fig.1.**
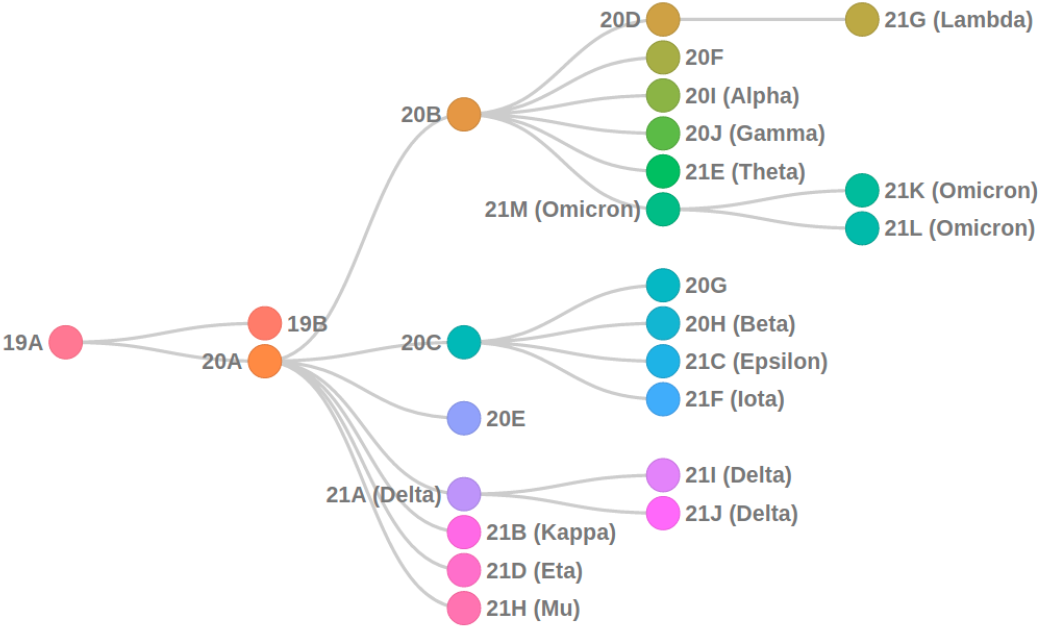
Nextstrain clades

### B. Mutations in spike protein

Spike protein, one of the structural proteins of SARS-CoV-2, spanning the genomic position from approximately 22,000 - 25,500, has caught significant attention because of the high frequency of mutations occurring in its region. A few of these mutations such as substitution N501Y in lineage B.1.1.7 (alpha variant) (5) and N439K and D614G in the variant 20A/S (3) are responsible for causing adverse effects on human health in multiple ways - increasing the affinity of the virus to bind to human cells or neutralising the effect of antibodies or increasing viral replication (6, 7). Therefore, to explore how such mutations are emerging through various clades over time and assess their impact on public health, computational methods are necessary. Due to its importance in the dynamics of mutations and their impact on human health, spike protein is chosen in this work to create datasets for developing and training an encoder-decoder neural network to model the evolution of protein sequences in adjacent clades of the phylogenetic tree (Figure 1). Only substitutions have been taken into account in our work as substitutions form the bulk of all the mutations occurring in most of the Nextstrain clades.

### C. Language translation and protein evolution

The field of language translation leverages the ability of the encoder-decoder neural network to translate sentences or paragraphs written in a source language (for example English) to a target language (for example German). Encoderdecoder neural network learns the latent representation from the source language and using this representation, it generates sentences in the target language after learning how to transfer from the source to the target language (8, 9). An analogy can be drawn from the language translation and learning evolution of genomic sequences as in both tasks, learning a model is required that knows how and what to change from one form of data to another. Building on this idea, the protein sequences are structured in a similar way to a language translation task - the source language resembles the protein sequences in the source clade (20A) and the target language resembles the sequences in the target clade (20B). The neural network learns the latent representation or edit mechanism to translate the sequences from the source clade to the target clade. The learned models can be used to generate sequences for unseen clades as the vocabulary of spike protein sequences are the same. However, this is not feasible for language translation tasks as vocabularies are not the same for different languages. Therefore, a language model trained in English-German languages cannot produce desirable outcomes in translating from English to French.

## Related work

Successful studies that implement encoder-decoder neural networks in learning transformation of the source language to the target language have motivated researchers to apply similar architecture for learning evolving biological sequences from one step to another. In (10), an autoencoder, a variant of an encoder-decoder neural network, is used to learn the latent representation of DNA sequences. In an autoencoder, source and target sequences remain the same. The encoder learns the latent representation of the source sequences and the decoder reproduces the same source sequences while minimising the reconstruction error. Following this approach, the authors achieved a splice site classification accuracy of 99% using the latent representation of DNA sequences learnt by the encoder that consists of bidirectional long-short term memory (LSTM) neural network layers. Variational autoencoder has been used in (11) to learn a continuous representation of SARS-CoV-2 protein sequences by minimising reconstruction error and the latent representation of protein sequences is used to produce their temporal variants using Gaussian processes. This approach uses the training data until May 30, 2020, which may lead to limited variability in sequences. Moreover, instead of learning the inherent amino acid distribution using weights to balance their occurrences in protein sequences, this approach achieves balancing by artificially increasing the number of protein sequences. MutaGAN in (1) uses a generative adversarial network in addition to an encoder-decoder network to simulate protein evolution in the influenza virus. The encoder-decoder network learns the editing mechanism of the source sequences to target sequences. Then, a discriminator, another neural network, distinguishes between pairs of real and fake pairs of source-target sequences to make the generation of generated sequences robust. Our approach takes into account the natural progression of protein sequences through two phylogenetically related clades (20A as the source and 20B as the target) to learn the transition mechanism as neural network models. These models, generate sequences that are closer to the ground truth protein sequences of clades that originate from the target clade 20B.

## Methods

In this approach, a method is proposed that jointly optimises an encoder-decoder neural network to learn and generate novel spike protein sequences. The encoder learns a latent representation of source protein sequences and the decoder uses this representation to translate the source protein sequences to the target protein sequence. The joint optimisation of encoder and decoder minimises the loss between the true target protein sequences and generated protein sequences. After the training is converged, the encoder-decoder neural network acquires the knowledge of how to translate source sequences to target sequences. Comparing the generated protein sequences to ground truth ones, substitutions at multiple genomic positions can be ascertained and further analysed.

### D. Data preparation

In this approach, SARS-CoV-2 spike protein sequences are used and they were collected from GISAID ^1^ (22–24). The FASTA file contains several protein sequences without clade information. Nextclade (2) tool in Galaxy (13) is used to assign a clade to each protein sequence. Using clades of protein sequences, pairs of sequences are created belonging to adjacent clades of the phylogenetic tree (shown in figure 1) of the evolution of SARS-CoV-2 sequences. Pairs of sequences that belong to adjacent clades such as 20A and 20B, exhibiting parent-child relationships, are created. It is assumed that sequences in 20A are predecessors to sequences in 20B based on the phylogeny. All versus all combinations of sequences are created using the sequences that belong to clades 20A and 20B. For example, 20 sequences in 20A and 30 in 20B will make 20 x 30 = 600 pairs of sequences. A few preprocessing steps are applied to these pairs of sequences to create a clean and good quality dataset that contains fewer outliers.

### E. Quality control

Data quality is improved following multiple steps. First, all the sequences containing any other letter other than 20 amino acids are removed. Second, all duplicate sequences from both the clades are separately removed to avoid duplicate pairs appearing in the dataset. Third, the pairs that are within the range of 1 - 10 levenshtein distance are kept and all pairs over this range are removed to restrict the divergence between parent-child sequences as protein sequences show a lower percentage of mutations compared to the length of the genome (3). Using pairs of sequences with large levenshtein distance may not be biologically relevant. Only sequences of length 1273 are used and the sequences that are longer or shorter than 1273 are discarded.

### F. KMER

After the quality control steps, sequences from both the chosen clades (20A and 20B) are represented as k-mers (19). K-mers of length 3 (k = 3) are created for all combinations of 20 amino acids that add up to 8000 (20 x 20 x 20) unique k-mers. For example, in the given partial protein sequence MFGFFVLLPLVSSQCVNLTTRT…, k-mers of length 3, also called 3-mers, are extracted as MFG, FGF, GFF,.....,…, TTR, TRT. Each protein sequence of length 1273 is converted to 3-mers resulting in a transformed sequence of length 1271. Each k-mer in these sequences is represented by an integer. All these sequences of k-mers are then converted into sequences of integers and each k-mer is represented by a unique integer. A collection of these sequences of integers for two clades are further divided into matrices, one matrix for each clade - the source clade (20A) and another for the target clade (20B). These matrices are split into training (80%) and test (20%) datasets. The training dataset is used in the encoder-decoder neural network for learning and the test dataset is used for evaluating trained models.

#### F.1. Encoder-decoder neural network

##### Network architecture

Encoder-decoder neural network for creating artificial protein sequences has two sub-networks, encoder and decoder, that are optimized jointly. High-level neural network architecture is shown in figure 2.

**Fig.2.**
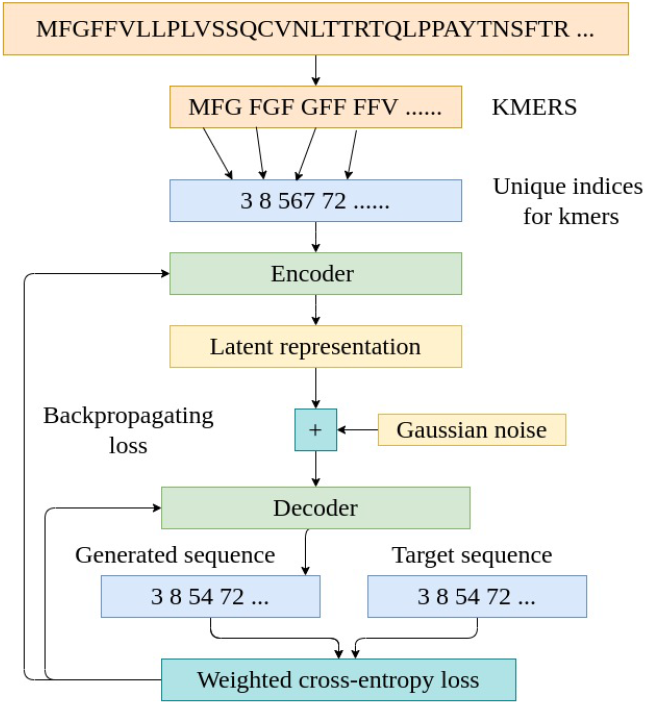
High-level neural network architecture

##### Encoder

The encoder network consists of an embedding layer and a bidirectional gated recurrent unit (GRU) layer, a variant of the recurrent neural network layer. The embedding layer creates a vector of fixed length for each integer that represents a k-mer. This vector becomes the embedding vector for the k-mer and is replaced by this vector during neural network learning. The second layer, GRU, is predominantly used for learning time-dependent data or sequential data. Each step of a sequence is dependent on previous steps that generalise to the connectivity of multiple steps. In this approach, we use protein sequences that are 1273 amino acids long. Therefore, long sequence modelling layers such as GRU are suitable that robustly learning the connections in such sequences ((20, 21)). In addition, GRU is wrapped around another layer, a bidirectional layer, and together they create a latent vector of fixed length for each protein sequence by reading them from both ends. The bidirectional GRU formulates two encoding vectors, one after reading a sequence forward and another backward. These two vectors are concatenated to make one encoding vector for each protein sequence that is further used by the decoder for a generating sequence.

##### Decoder

The decoder has 3 different types of layers. The first one is an embedding layer that is similar to the encoder. As in the encoder, it is connected to the GRU layer. The concatenated vector from the encoder is used to initialise the GRU layer of the decoder. In contrast to the encoder, the bidirectional layer is not used in the decoder as the generation of target sequences and computation of latent representation happen at each genomic position and not for the entire sequence at once. This technique for generating output at each position is known as teacher forcing and it helps the neural network learn the structure of sequences faster (14). Using this technique, the generation of protein sequence happens at each genomic position and then all generated amino acids are combined to produce an entire protein sequence. At each genomic position, the true output, an amino acid from the true target sequence, is fed back to the decoder along with its latent representation computed at the previous step. Together, they generate the most probable amino acid for the next position and the latent representation is also updated. The output of the GRU layer is then fed to a dense layer with softmax activation. An argmax function is applied to the logits generated by this dense layer to create a sequence of integers that is essentially a sequence of k-mers. The task of generating sequences is reduced to a classification task where, at each genomic position, an integer that belongs to an amino acid is predicted. Suitable dropout layers are applied between each pair of layers in both, encoder and decoder networks to prevent overfitting.

##### Class balancing

The dataset consisting of pairs of source and target sequences suffers from significant imbalance at many genomic positions. At each genomic position, for most of the pairs, one amino acid is located and substitution occurs only at a few positions. When the encoder-decoder network is trained on such a dataset without implementing any approach to balance the occurrences of amino acids at different genomic positions, the network learns to generate only the dominant amino acid at each position while ignoring different amino acids that are valid substitutions. Therefore, it generates the same target protein sequences irrespective of the presence of multiple variants of source protein sequences. To mitigate such data imbalance, an approach from (15) is implemented that works nicely with protein sequences as well. The approach approximates, in equation 1, an expected number of samples for each class. This expected number of samples for each class (an amino acid) is a real number and is multiplied by the cross-entropy error between the true and generated set of amino acids at each genomic position as the sample weight. Samples that belong to the less dominant amino acids are weighted strongly and samples belonging to more dominant amino acids are weighted weakly. Multiplication of these weights to the loss function enables the neural network to focus on less dominant amino acids too as their misclassification would amount to larger loss compared to the misclassification of more dominant amino acids because of larger weights of less dominant amino acids. In this approach, the frequency of all substitutions present in the training data is computed at each genomic position. Using the frequencies of substitutions at each genomic position, the expected number of samples for each amino acid is computed. Then, the weights are computed using equation 1 and then multiplied by the cross-entropy loss between the true and generated set of amino acids. Using this updated loss function, the generated target protein sequences show similar variations at most of the genomic positions as shown in figure 4. In equation 1, *β* is a hyperparameter. The number of samples for each amino acid is represented by *n_y*.

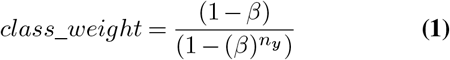

##### Stateful GRU

Recurrent neural network (RNN) layers such as GRU in a neural network are good at learning connections between multiple temporal steps. But, when sequential data contains hundreds of temporal steps, the performance of GRU in predicting future connections (at future time-steps) suffers as it becomes hard to maintain memory for such long sequences without losing important information. To make learning on sequential data with hundreds of temporal steps, stateful recurrent units are used (16, Steven Elsworth and Stefan Güttel 2020). To enable stateful training with GRU, each set of source and target protein sequences are divided into batches, known as stateful batches, of fixed length. The first divided set of sequences is used for learning the second divided set of sequences, the second is then used for the third and so on. By using this, GRU only needs to maintain the memory of only the fixed length of stateful batches at a time and not for the entire length of protein sequences. For example, a protein sequence of length 100 is divided into 2 stateful batches of length 50. With this division, GRU needs to remember states of only 50 amino acid long sequences. The first stateful batch of 50 amino acids is used to predict the next stateful batch of 50.

#### F.2. Experiments

Quality control steps applied on the pairs of source-target spike protein sequences make approximately 10,000 pairs of good quality. 80% of these pairs are used for training the neural network and the remaining 20% for evaluating it. Training and test datasets contain two columns, one each for sequences in 20A and the second for 20B. The sequences in 20A are used to generate latent representations of source sequences by the encoder. In the data preparation process, all versus all combination is used which leads to cases where each sequence in 20A gets mapped to multiple sequences in 20B. Therefore, the latent representations of each sequence in 20A is added to a Gaussian distributed noise with zero mean and unit standard deviation to incorporate small variations in each of the latent representations as they are used by the decoder to generate slightly different sequences of 20B. These latent representations are used to initialise the GRU layer of the decoder. The decoder consumes the latent representations and target sequence’s amino acids, following the approach of teacher forcing, and generates amino acids for each genomic position. For generating amino acid at the first genomic position, a list of 0s is passed as the initialisation to the decoder. The neural network is trained for 6 epochs (6 times on the entire set of training pairs of sequences) and at each epoch, the model is saved. These models are used to generate multiple variations of sequences using 20A sequences in the test dataset.

##### Hyperparameters

The encoder-decoder neural network has several hyperparameters such as the latent size of encoded vectors (GRU units)/latent representation, the number of samples for each training iteration (batch size), length of sequence for stateful training (stateful size), embedding dimension for each kmer and learning of the adam optimiser. The number of GRU units is set to 256 for learning the entire length of the sequence creating a vector of dimension 512 for each sequence. The higher it is, the stronger is the model. The batch size is set to 8. The stateful size is set to 41 creating 31 batches of each sequence of length 1271. This number is chosen so that it creates stateful batches of equal size. Each batch provides the initial latent representation for generating the next batch. The embedding dimension is set to 128 fixing the size of the vector with which each k-mer is represented. The k-mer length is chosen as 3 and not higher because increasing the k-mer size would result in a higher number of possible combinations of 20 amino acids. It will increase the matrix size that contains the generated sequences, in the form of logits, and will require more memory. The learning rate is chosen as 0.01 for faster convergence of the neural network for the adam optimiser. The error function is the cross-entropy loss weighted by the expected number of samples for each amino acid at each genomic position and is computed using equation 1. The dropout rate is set to 0.2 to prevent overfitting. Beta in equation 1 is set to 0.9999 which computes weights for each amino acid proportional to its expected occurrence in the training data.

##### Softwares

Tensorflow 2.7.0 is used for creating the architecture of the encoder-decoder neural network using Python 3.9.7. Bio 1.79 is used to read FASTA files containing spike protein sequences. For faster training, 1 Nvidia Tesla T4 GPU is used running on CUDA 11.4. The model needs around 1 day for 1 iteration.

## Results

The neural network was trained for 6 iterations over the entire training dataset of approximately 10,000 pairs of protein sequences. After analysing the generated sequences using models trained at different iterations of training, it is found that different models tend to generate variations at different genomic positions. To allow the generated sequences to exhibit variations at multiple genomic positions where the training data also shows variations, an ensemble of models is used to generate sequences. Sequences from clade 20A in the test dataset are used to generate sequences. While generating, the levenshtein distance between each source sequence of 20A and its corresponding generated sequence is calculated. Pairs of source-generated sequences are discarded if the levenshtein distance is greater than 10 and those pairs within the levenshtein distance of 1-10 are stored. Out of 4800 generated sequences, 3932 (82%) pairs of source-generated sequences are within the levenshtein distance range of 1-10. To find out how the amino acids get substituted from the source to target clades (20A to 20B), a heatmap (figure 3) is created that shows the transformation. On the y-axis, the frequency of substitution of each amino acid in 20A sequences to each amino acid in 20B sequences is shown for training (A) and generated (B) (source sequences in 20A from the test dataset) datasets. It is evident from the strong pearson correlation (> 0.7) between A and B that the generated sequences resemble the ground truth sequences in clade 20B. The frequencies are normalized by their respective dataset sizes.

**Fig. 3.**
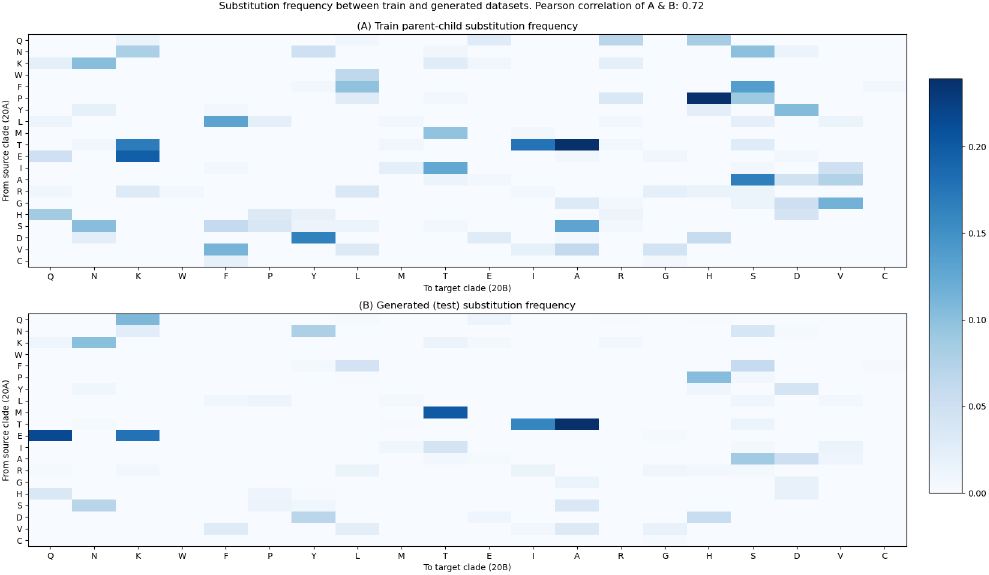
Protein substitutions in true and generated sequences

Figure 4 shows the frequency of substitutions at each genomic position for the sequences of length 1273 in training (A) and generated (B) datasets. Each frequency of substitution is normalized between 0 and 1 to make the bar plots comparable. The frequency of substitutions shown in Figure 4 (A) is calculated using the sequences in clades 20A (as the source) and 20B (as the target) from the training dataset. These are the only substitutions that are presented to the proposed encode-decoder neural network during training. Figure 4 (B) shows the substitution frequency between the ground truth sequences of clade 20A in the test dataset and the sequences that are generated using these ground truth sequences of clade 20A in the test dataset. There is a high pearson correlation (>0.7) between figures (A) and (B) showing that the generated sequences have a good resemblance with the ground truth sequences in clade 20B. Genomic regions of the high variability of the entire length of spike protein are evident in both the bar plots and they show resemblance not only for genomic positions that are substituted with high frequencies (high peaks) such as regions around 480 and 680 but also for those that are substituted with low frequencies (low peaks) such as 81 and 210. Exhibiting such a combination of high- and low-frequency substitutions brings them closer to naturally occurring spike protein sequences. Moreover, it also validates that the deep learning models learn the capacity to generate more realistic sequences that have a good similarity to the natural distribution of substitutions in adjacent clades, 20A and 20B. These models are used to generate sequences that occurred in later clades such as the children clades of 20B of the phylogenetic tree (shown in figure 1). These novel sequences are then compared to the ground truth sequences that belong to the children clades of 20B.

**Fig. 4.**
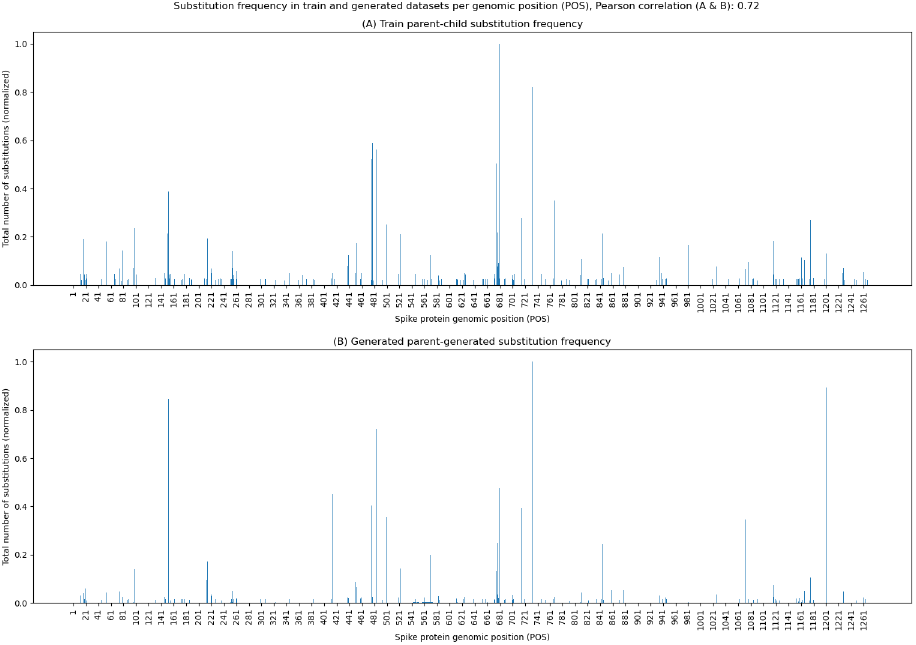
Protein substitutions distribution in true and generated sequences (20A-20B-generated)

### G. Per position substitution distribution

Figure 5 shows a similar plot as in Figure 4, but in this figure, frequencies of substitutions are calculated using the reference spike protein sequence (Wu, F. *et al.* 2020). Figure 5 (A) shows the frequency of substitutions between the reference spike protein sequence and the ground truth sequences of 20B in the training dataset and figure 5 (B) shows the frequency of substitutions between the reference spike protein sequence and the sequences generated using sequences in 20A from the test dataset. Comparing these two bar plots, it can be seen that many high peaks of substitutions are robustly captured in the generated sequences. For example, the substitution D614G is strongly captured in the generated sequences which are also present in figure 5 (A). Comparing figure 5 to figure 4 (A), it is found that the neural network train on a little activity at position 614 but robustly generates sequences that contain high substitutions at the same position (D614G). Moreover, multiple genomic regions of high variability such as near genomic positions 480, 680 and 730 in both training and generated datasets show high similarities. These examples show that the generated dataset closely resembles the ground truth sequences that belong to clade 20B. It suggests that the neural network captures latent features from sequences in clades 20A and 20B and generates such sequences that contain substitutions in a similar distribution as found in naturally occurring spike protein sequences. Moreover, a high pearson correlation of > 0.8 also suggests that there exists a strong relationship between ground truth sequences of clade 20B and the generated sequences.

**Fig. 5.**
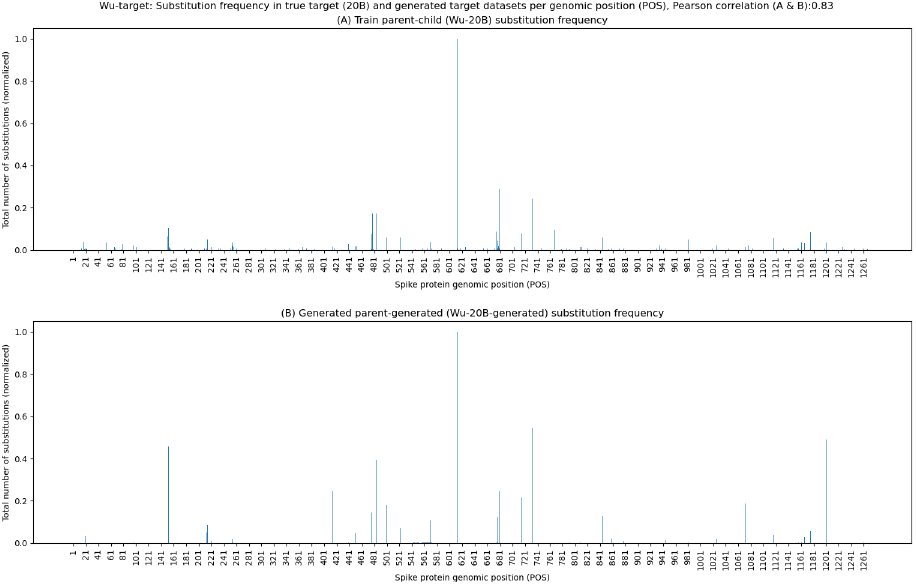
Protein substitutions distribution in true and generated sequences

### H. Predicted substitutions in future clade (20J)

Figure 6 shows a similar plot showing substitution frequencies as in figure 5 using the same neural network models but using different source sequences. Figure 6 (A) shows substitution frequencies between reference spike protein sequences and sequences that belong to clade 20J (Gamma, see figure 1). Clade 20J (Gamma) is one of the children clades of 20B. Figure 6 (B) shows substitution frequencies between reference spike protein sequences and sequences that are generated using the ground truth sequences of 20B from the test dataset. Note that for figure 5 (B), 20A sequences from the test dataset are used. It can be seen in the plot that the distribution of substitution frequencies in figures 5 (B) and 6 (B) are slightly different even though the used trained models for generation are the same. The only difference is the set of source sequences that are used for the generation, for figure 5 (B), test sequences that belong to 20A are used and for figure 6 (B) test sequences that belong to 20B are used. For example, comparing figures 5 and 6, it is seen that there is a small number of true and generated substitutions at position 1176, which is a defining substitution for clade 20J (Gamma) (**?**), in figure 5. But, the number of substitutions at position 1176 increases in figure 6, both for true and generated bar plots. In addition, the substitution shows a higher activity for the generated sequences (figure 6 (B)) at multiple other locations such as 484, 501 and 614 similar to the pattern shown by the ground truth 20J (Gamma) sequences. In a separate task, the same set of trained models, trained on pairs of sequences from 20A and 20B, is used to generate sequences for a different branch, originating from clade 20C, of the phylogenetic tree of clades. But, unfortunately, the outcome of such a generation task is not encouraging which shows a limitation of these models.

**Fig. 6.**
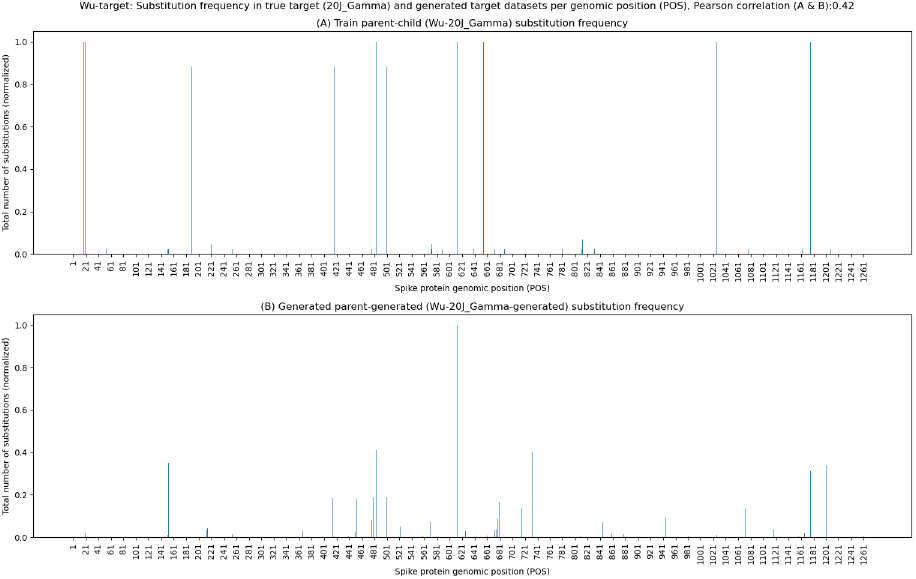
Protein substitutions distribution in true (Wu-20B) and generated sequences (Wu-20J)

### I. Alphafold

One of the generated sequences by the ensemble of trained models is used to generate its 3D structures using Deepmind’s AlphaFold as shown in figure 7. Most of the regions of the 3D structure of the generated spike protein show high confidence (denoted by blue colour) and only a few have low confidence (denoted by red colour). It lends evidence to the fact that the generated protein sequence by deep learning models has high a resemblance to a naturally occurring spike protein of the SARS-CoV-2 virus (18).

**Fig. 7.**
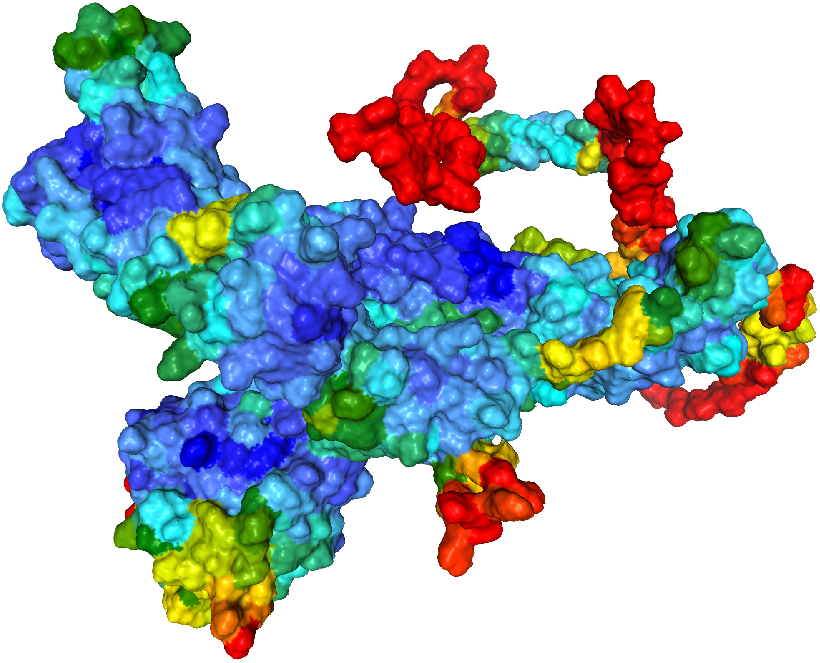
Alphafold generated structure of a generated protein sequence

## Discussion

The proposed approach makes use of deep learning to learn the evolution of spike protein sequences through pairs of sequences, each belonging to one clade. The neural network learns the transition between pairs of sequences and an ensemble of trained models is used to generate spike protein sequences that show resemblance to spike protein sequences that originate from target clade 20B. When comparing the generated spike protein sequences with the original reference sequences, the resemblance of distribution (person correlation > 0.7), at a few genomic locations, between ground truth sequences and generated sequences is encouraging and shows potential for further research. It is feasible to use this approach to learn from the already evolved sequences to artificially create spike protein sequences that have not yet been evolved and get insights into the genomic positions that indicate higher activity. In addition, the approach develops a general algorithm that can be used with other structural and non-structural protein sequences associated with SARS-CoV-2 and with other living beings too. But, there are a few limitations of this approach. One is related to data preparation. The sequences of clades 20A and 20B have been chosen randomly from a pool of clade labelled sequences. The clade 20A came before 20B, therefore, it is important to choose sequences of clade 20A that have been collected before sequences of 20B. The categorisation of protein sequences into clades by the Nextclade tool may involve errors. Moreover, there is an unequal distribution of the submitted sequences based on geography. Around half of the submitted sequences for clades 20A and 20B in GISAID come from the US and the rest of the nations contribute to the other half of these clades. In addition to mitigating such biases in the training data, it is also important to train models on different branches of the phylogenetic tree and compare their performances.

## Author Approvals

All authors have seen and approved the manuscript, and it hasn’t been accepted or published elsewhere.

## Acknowledgements

We gratefully acknowledge all data contributors, i.e. the Authors and their Originating laboratories responsible for obtaining the specimens, and their Submitting laboratories for generating the genetic sequence and metadata and sharing via the GISAID ^2^ Initiative (22–24), on which this research is based.

## Supplementary information

Codebase of this project is available here: online.

---

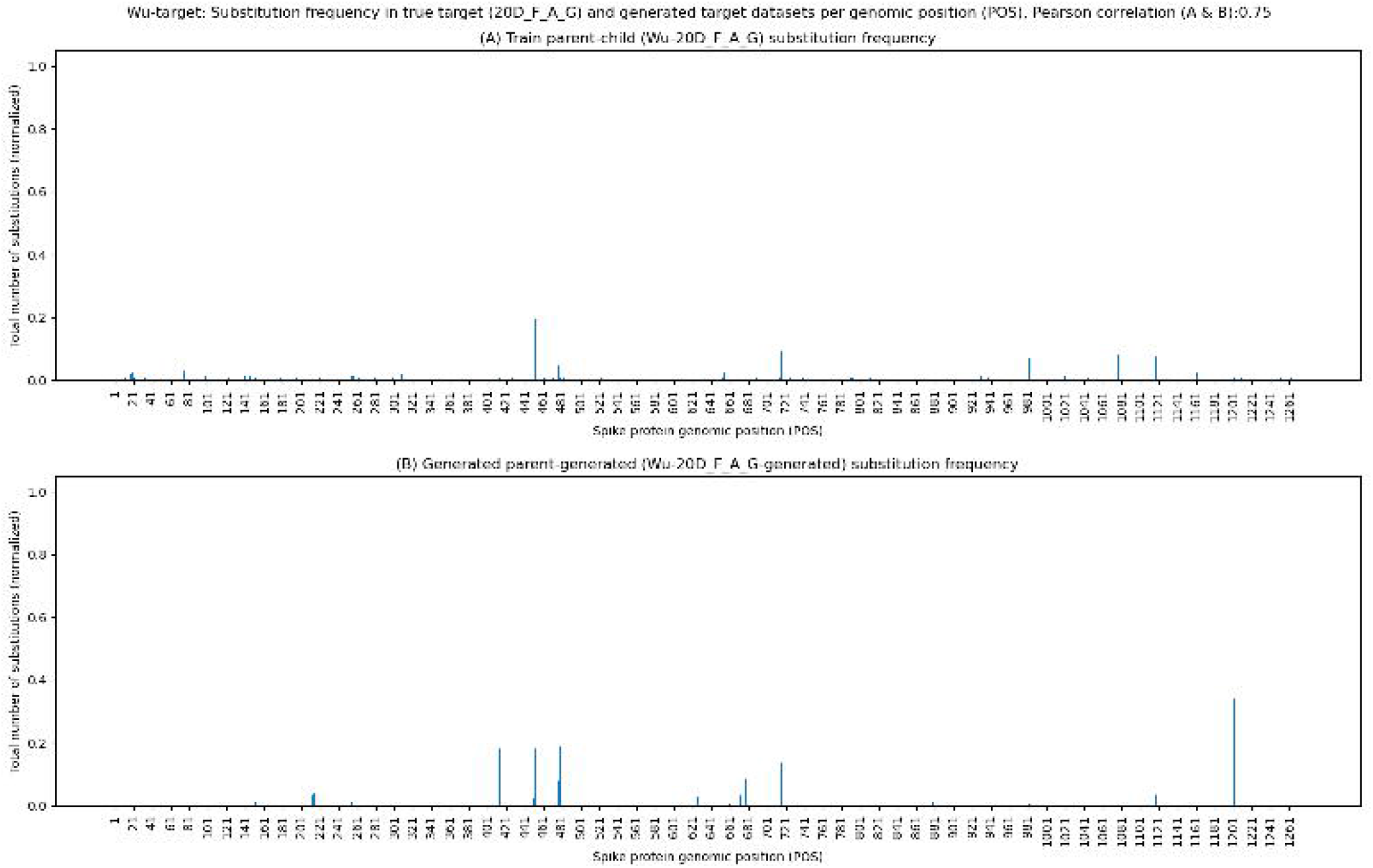

## Footnotes

1 EPI_SET_20220316ph

2 EPI_SET_20220316ph

